# Competitiveness prediction for nodule colonization in *Sinorhizobium meliloti* through combined *in vitro* tagged strain characterization and genome-wide association analysis

**DOI:** 10.1101/2020.09.15.298034

**Authors:** A. Bellabarba, G. Bacci, F. Decorosi, E. Aun, E. Azzarello, M. Remm, L. Giovannetti, C. Viti, A. Mengoni, F. Pini

**Affiliations:** Department of Agronomy, Food, Environmental and Forestry (DAGRI), University of Florence, Via della Lastruccia 10, I-50019, Sesto Fiorentino (Italy); Genexpress Laboratory, Department of Agronomy, Food, Environmental and Forestry (DAGRI), University of Florence, Via della Lastruccia 14, I-50019, Sesto Fiorentino, Italy; Department of Biology, University of Florence, Via Madonna del Piano 6, I-50019 Sesto Fiorentino, Italy; Department of Bioinformatics, Institute of Molecular and Cell Biology, University of Tartu, Riia 23, 51010 Tartu, Estonia; Department of Biology, University of Bari Aldo Moro, via Orabona 4, I-70124 Bari, Italy

**Author notes:** **Corresponding authors** Alessio Mengoni, Department of Biology, University of Florence, via Madonna del Piano 6, 50019, Sesto Fiorentino, Italy., Carlo Viti, Department of Agronomy, Food, Environmental and Forestry (DAGRI), University of Florence, Via della Lastruccia 10, I-50019, Sesto Fiorentino, (Italy).

**Keywords:** GWAS, competition, *Sinorhizobium meliloti*, rhizobia, legume

## Abstract

Associations between leguminous plants and symbiotic nitrogen fixing bacteria (rhizobia) are a classical example of mutualism between a eukaryotic host and a specific group of prokaryotic microbes. Though being in part species-specific, different strains may colonize the same plant symbiotic structure (nodule). It is known that some rhizobial strains are better competitor than others, but detailed analyses aimed to predict from the rhizobial genome its competitive abilities are still scarce. Here we performed a bacterial genome wide association (GWAS) analysis to define the genomic determinants related to the competitive capabilities in the model rhizobial species *Sinorhizobium meliloti*. Thirteen tester strains were GFP-tagged and assayed against three reference competitor strains RFP-tagged (Rm1021, AK83 and BL225C) in a *Medicago sativa* nodule occupancy test. Competition data in combination with strains genomic sequences were used to build-up a model for GWAS based on k-mers. The model was then trained and applied for competition capabilities prediction. The model was able to well predict the competition abilities against two partners, BL225C, Rm1021 with coefficient of determination of 0.96 and 0.84, respectively. Four strains showing the highest competition phenotypes (> 60% single strain nodule occupancy; GR4, KH35c, KH46 and SM11) *versus* BL225C were used to identify k-mers associated with competition. The k-mers with highest scores mapped on the symbiosis-related megaplasmid pSymA and on genes coding for transporters, proteins involved in the biosynthesis of cofactors and proteins related to metabolism (i.e. glycerol, fatty acids) suggesting that competition abilities reside in multiple genetic determinants comprising several cellular components.

**IMPORTANCE:** Decoding the competitive pattern that occurs in the rhizosphere is challenging in the study of bacterial social interaction strategies. To date, single-gene approach has been mainly used to uncover the bases of nodulation, but there is still a gap about the main features that *a priori* turn out rhizobial strains able to outcompete indigenous rhizobia. Therefore, tracking down which traits make different rhizobial strains able to win the competition for plant infection over other indigenous rhizobia will allow ameliorating strain selection and consequently plant yield in sustainable agricultural production systems. We have proven that a k-mer based GWAS approach can effectively predict the competition abilities of a panel of strains, which were analyzed for their plant tissue occupancy by using double fluorescent labeling. The reported strategy could be used for detailed studies on the genomic aspects of the evolution of bacterial symbiosis and for an extensive evaluation of rhizobial inoculants.

## INTRODUCTION

The nitrogen-fixing symbiotic interaction is a classic example of a mutualistic association (1). The interaction between rhizobia and host plants (mostly *Fabaceae*, but also with the genus *Parasponia*) starts with an exchange of molecular signals: flavonoids released from the plant roots and Nod factors produced by rhizobia (2). Nod factors induce a molecular pathway on plant root cells, which ultimately leads to rhizobia entry in the tissue and intracellular colonization. Intracellular rhizobia differentiate into bacteroids, which express the nitrogenase, the enzyme responsible for the fixation of atmospheric di-nitrogen to ammonia (3, 4). On the plant root, a new structure, called nodule, forms, where rhizobia can proliferate and differentiate into bacteroids (5, 6). Indeed, in a single nodule (a mass of a few hundreds of milligrams), up to 10^6^ bacterial cells can be recovered, while in the soil, free-living rhizobia are usually not more than 10^3^-10^4^/g of soil (7).

As in a trade framework, the benefit for the rhizobium is a protected environment where it can proliferate (under control) and receive carbon and energy supply from the plant, while the reward for the plant is the availability of fixed nitrogen (8, 9). Rhizobia transmission is horizontal among plants, and plants may be colonized by poorly effective (low nitrogen fixing) strains. However, host plants could control the spread of inefficient strains by sanctioning root nodules, hence limiting the growth of underperforming strains (10–12). Moreover, the presence of multiple strains within a nodule (mixed nodule) may occur (7, 13). Under these circumstances, inefficient strains can behave as cheaters decreasing the overall nitrogen-fixing performance (7, 8). Consequently, understanding the competition between strains has a great importance either to study the evolution of symbiosis (14, 15) and to predict how much efficient would be a rhizobial inoculant for agricultural applications (16). The genetic bases of competitiveness among rhizobial strains are still elusive and, most of the studies, addressed symbiotic genetic determinants from experiments involving mutants of a few different natural strains (7).

The link between a phenotype and its genetic basis, hence predicting phenotypes from the sole genomic information is one of the great challenges of biology (17). Genome-Wide Association Studies (GWAS) are commonly used for identifying the putative functional role of set of allelic variation in groups of individuals. In bacteria, GWAS have been applied to several species for predicting complex (i.e. multigenic) phenotypes, as antibiotic tolerance and host interaction (see for examples (18–20)). However, most studies were related to phenotypes under strong selective pressure, while it is still challenging to determine the genetic basis of phenotypes under mild selection (21). The identification of the genetic determinants in host-bacteria interaction is essential, and GWASs on plant holobionts (the ensemble of the plant and the other organisms living in or around) have been proposed (21), aiming to provide the basis for future breeding programs, which includes among the plant trait, the recruitment of the “good” microbiome.

Recent studies have reported the feasibility of experimental setups combining symbiotic assays with genome sequencing approaches and GWAS in rhizobia to define the genetic determinants of symbiotic performances (24–27). In the model rhizobium *Sinorhizobium meliloti*, association analysis has been used to explore the genetic basis of various phenotypic traits including antibiotic resistance, symbiotic and metabolic traits (24). A select and re-sequence approach has been successfully applied to measure the fitness of a set of 101 *S. meliloti* strains against two genotypes of the host plant *Medicago truncatula* (25). However, the predictive value of single rhizobial genotypes (i.e. genomes) toward the expected fitness in terms of competitive capabilities to establish successful symbiosis is still unclear. Indeed, to date, though many genetic details of the symbiotic interaction are known for single strain colonization (28), there is still a gap about which rhizobial features increase the chances to win the competition for plant infection outcompeting others indigenous rhizobia. Therefore, unearthing these genetic determinants can promote studies on the genomic aspects of the evolution of bacterial symbiosis and can have direct practical application in the screening and amelioration of rhizobial inoculants to be used in sustainable agricultural production systems.

Here, we aimed to address the possibility to predict, on the basis of the sole genome sequence, the competitive capabilities of rhizobial strains. We used as model rhizobial species, *Ensifer* (syn. *Sinorhizobium*) *meliloti* for which many molecular genetics data and tools are available (29), a good number of strains has been sequenced and preliminary data on symbiotic performances and competition are present (7, 18, 19, 25, 30). We settled a series of experiments where pairs of fluorescently labelled *S. meliloti* strains were used to infect alfalfa (*Medicago sativa*) plants. The measure of the competition phenotypes of each strain was then used with genomic sequences of the same strain to perform genome-wide association analysis and construct a predictive model of competition based on genomic features (k-mers) occurrence.

## RESULTS

### Construction of fluorescently tagged *S. meliloti* strains

To set up *in vitro* tests for measuring competition capabilities, a panel of 16 *S. meliloti* strains was selected. Three well characterized strains for competition capabilities (*S. meliloti* BL225C, AK83 and Rm1021) were chosen (7) and used as reference competitors against 13 *S. meliloti* strains (tester strains) whose genome sequences were available (Table S1). Phylogenetic relationships among the 13 *S. meliloti* strains were evaluated (Fig. S1A) and their pangenome was analyzed (Fig. S1B-D). The pangenome is composed of 15419 genes: 4278 are shared by all strains (core genome) while 6622 genes are strain specific (Fig. S1D). For all the above-mentioned 13 tester strains, GFP (green fluorescent protein) derivatives were constructed by cloning the pHC60 plasmid, which constitutively expresses the GFP. *S. meliloti* Rm1021, BL225C and AK83 strains were tagged with RFP (red fluorescent protein) by using the pBHR mRFP plasmid. Preliminary single inoculation assays were performed showing that all strains were able to form nodules on the root of *M. sativa* plants (Fig. S2). For all but two strains (M270 and T073) nitrogenase activity inside nodules was detected (Fig. S2D), in agreement with previous results that showed low nitrogen fixation abilities in symbiotic interaction with *M. truncatula* (31). Differences in nodulation, plant growth and nitrogenase activity among strains were observed. *S. meliloti* AK58, RU11/001, SM11, USDA1157, GR4, and CCMM B554 strains gave the highest values of nitrogenase activity and plant growth promotion (Fig. S2).

### Competition capabilities for colonization of nodules differ in relation to competitor counterpart

Tagged *S. meliloti* strains were used in a set of competition experiments: each with a GFP-tagged strain (13 strains in total) *versus* an RFP-tagged competitor strain (*S. meliloti* BL225C, AK83 or Rm1021) (13 x 3 total of 39). Large variability in nodule colonization was observed among and within the three sets of competitions experiments (*vs* Rm1021, *vs* AK83 and *vs* BL225C). The three competitions test showed differences in the number of total nodules produced *per* plant (p < 0.001, Fig. S3A) and, in the competition experiments with AK83 the highest number of nodules was observed (Fig. S3A).

Competition capabilities were evaluated as single nodule occupancy (nodules colonized by a single strain) of the tested strain in respect to a reference strain; good competitors were characterized by a single nodule occupancy higher than 60%, medium competitor between 20 and 60% and weak competitors below 20%. In the competition with Rm1021, most of the strains outperformed (Fig. 1A). This competition test was characterized by a high average value of nodule occupancy of the strains tested equal to 65.12%. Most of the strains showed single nodule occupancy higher than 60%, with the highest value (93.4%) observed for GR4, three strains (HM006, Rm41 and M270) exhibited medium competition capabilities and two strains (T073 and USDA1157) lowly performed (Fig. 1A and Table S2). Among the three strains with medium competition capabilities, HM006 and Rm41 displayed values close to high performing strains (58.6% and 50.5%, respectively) while M270 nodule occupancy was 37%. Medium-low performance of strains M270 and T073 (T073 competition was characterized by mixed nodules or with strain Rm1021 only) may be related to nodule sanctioning (plant limiting nutrient to inefficient nodules) as T073 and M270 were unable to fix nitrogen (Fig. S2D). Concerning USDA1157 strain, which showed good nitrogen-fixing abilities, we might hypothesize that the lower value of nodule occupancy could be related to direct strain-by-strain competitive interaction, more than to plant sanctions.

**Figure 1.**
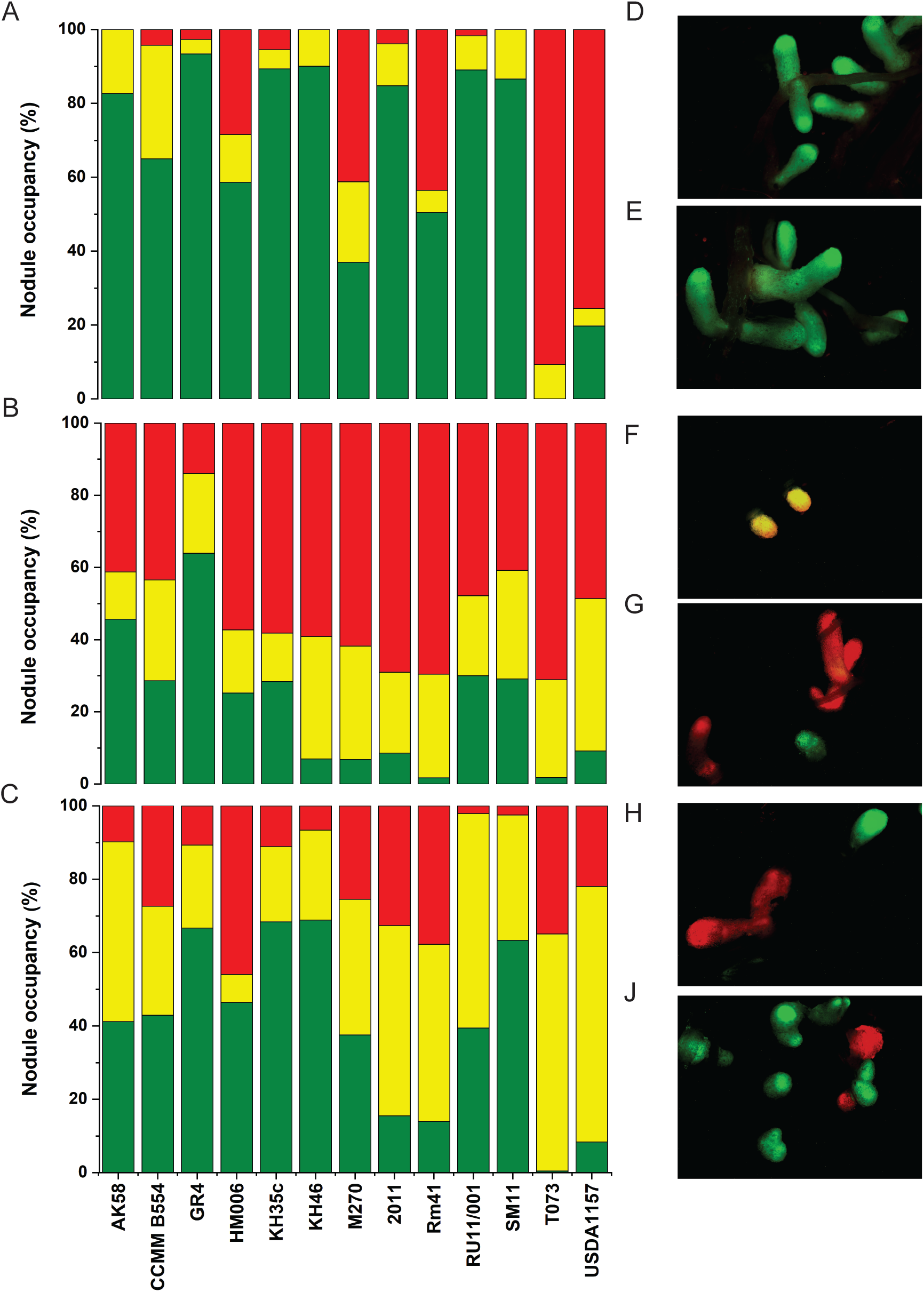
Competition performances and epifluorescence stereomicroscope images. Bar plots showing the percentage of nodule occupancy of 13 *S. meliloti* strains in three sets of competition experiments: A) competition against *S. meliloti* Rm1021, B) competition against *S. meliloti* AK83 and C) competition against *S. meliloti* BL225C. Green bars represent single nodule occupancy of the strains tested whose ID is reported on the x-axis, in yellow is showed the percentage of mixed nodules (nodules occupied by both strains) and in red the single nodule occupancy of the competitor used in each set of experiments. Pictures show nodules of *M. sativa* inoculated with a mix of *S. meliloti* 1021 RFP-tagged and D) KH46 GFP-tagged or E) GR4 GFP-tagged; F) and H) *S. meliloti* AK83 RFP-tagged and HM006 GFP-tagged; and *S. meliloti* BL225C RFP-tagged and I) CCMM B554 GFP-tagged or J) RU11-001 GFP-tagged.

In the competition experiments with AK83, conversely to the pattern highlighted when Rm1021 was used as a competitor, a general decrease of nodule occupancy of the 13 strains tested was observed (Fig. 1B), resulting in a lower average value of nodule occupancy (21.99%; Fig. 1B and Table S2). Except for GR4, that showed the highest percentage of occupancy equal to 63.9%, all strains displayed weak-medium competitive capabilities (nodule occupancy lower than 60%; Fig. 1B and Table S2). The lowest value of nodule occupancy equal to 1.7% and 1.8% were detected for Rm41 and T073, respectively.

An intermediate scenario between those above described was detected for the competition with BL225C (Fig. 1C and Table S2), where the average value of nodule occupancy of the strains tested was equal to 39.45%. The most competitive strains were GR4, KH35c, KH46, and SM11, showing a nodule occupancy ranging from 63.4 % to 68.9%. The lowest percentage of nodule occupancy equal to 0.4% was detected for T073.

In both the competition with AK83 and BL225C, a higher abundance of mixed nodules (nodules infected by both *S. meliloti* strains) was observed compared to the competition with Rm1021 (Fig. S3B).

### Modeling competition pattern from genome sequences

Short DNA oligomers with constant length k, termed k-mers, allow to capture a large set of genetic variants in a population, including SNPs, insertions/deletions (INDELs) (20, 32). To pinpoint genetic determinants that might be responsible for the competition capabilities variation among *S. meliloti* strains, we identified competitive phenotype-specific k-mers and trained a k-mer-based statistical model for predicting phenotypes of interest.

A matrix based on the competition phenotype (Table S2) and the genome sequences of the strains tested, was used to identify the most strongly associated-phenotype k-mers (p-value < 0.05) and linear regression models were built for each set of competition experiments. To measure the accuracy of predictive models, the strains were randomly split into a training test and a test set (32). The effectiveness of linear regression models in describing the three observed competition patterns varied among the dataset. The prediction was greatly accurate for the competition phenotype *versus* BL225C (Fig. 2C), as indicated by the coefficient of determination R^2^ equal to 0.98 and 0.96 in training and test set respectively (Table 1), and by the degree of similarity between actual and predicted phenotype (Fig. 2), The number of total k-mers identified in tested strains as significantly associated with competition against BL225C was smaller in comparison with other datasets (Table 1). The wider p-value range for significantly associated k-mers was founded in the analysis of competition against BL225C.

**Figure 2.**
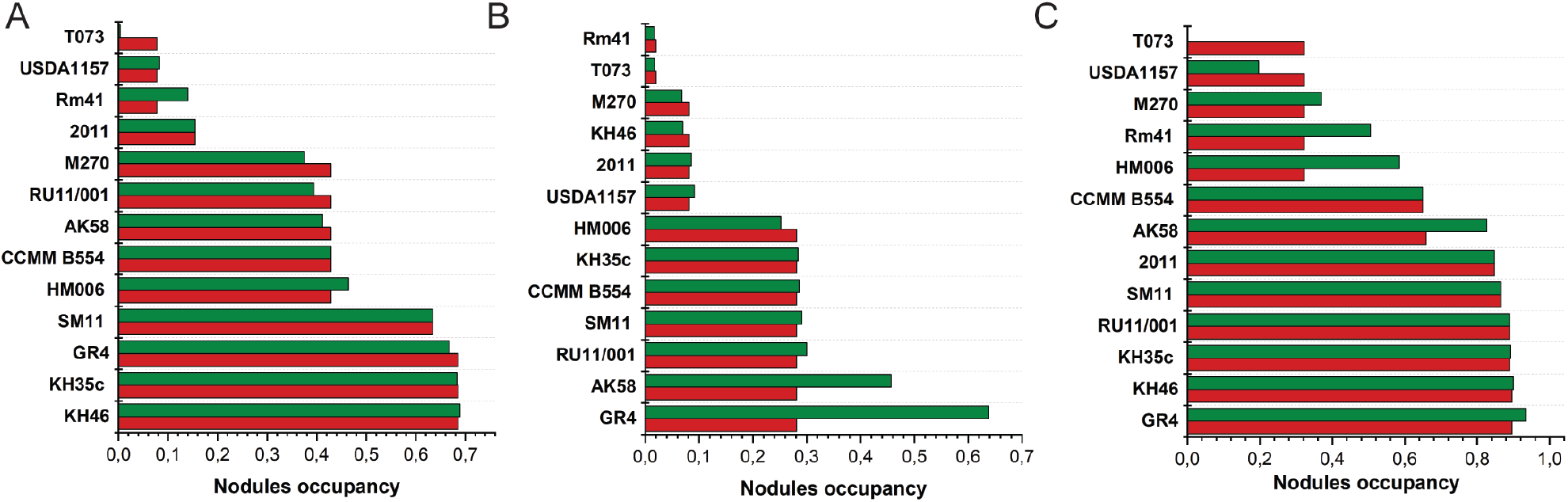
Predicted and actual competition patterns. Single nodule occupancy of *S. meliloti* strains in competition experiments versus *S. meliloti* strains A) Rm1021, B), AK83 and C) BL225C. Green bars indicate the actual phenotypes, red bars the predicted phenotypes.

**Table 1.** Performances of linear regression models for the three competition experiments. The results of both training and test set by PhenotypeSeeker are reported.

| Dataset |  | The mean squared error | The coefficient of determination (R<sup>2</sup>) | The Pearson correlation and p-value | The Spearman correlation coefficient and p-value | Total K-mers (p-value < 0.05) | Range p-value |
| --- | --- | --- | --- | --- | --- | --- | --- |
| Vs Rm1021 | Training set | 0.02 | 0.74 | 0.86, 0.0 | 0.98, 0.0 | 439886 | 4.99E-0<sup>2</sup> – 4.31E-0<sup>3</sup> |
|  | Test set | 0.01 | 0.84 | 0.96, 0.04 | 1.0, 0.0 |  |  |
| Vs BL225C | Training set | 0.0 | 0.98 | 0.99, 0.0 | 0.97, 0.0 | 182804 | 4.98E-0<sup>2</sup> – 1.35E-0<sup>5</sup> |
|  | Test set | 0.0 | 0.96 | 0.99, 0.01 | 0.89, 0.11 |  |  |
| Vs AK83 | Training set | 0.0 | 0.99 | 0.99, 0.0 | 0.92, 0.0 | 292884 | 4.97E-0<sup>2</sup> – 1.05E-0<sup>3</sup> |
|  | Test set | 0.04 | 0.11 | 0.8, 0.2 | 0.77, 0.23 |  |  |

In terms of model-evaluation metrics, the linear regression model was less reliable in defining the competition against Rm1021 for both training and test sets respect to the competition against BL225C (Table 1). In particular, the predictive capacity of the model was inaccurate for those strains that displayed low or medium-low competition capabilities (*S. meliloti* T073, USDA1157, M270, Rm41 and HM006) in a competition where most of the strains showed medium-high capacities (Fig. 2). Furthermore, due to the greater extent of positive performances showed by most of the strains tested against Rm1021, a higher amount of total k-mers related to this competition phenotype was obtained (Table 1). Concerning the level of accuracy of the predictive model for competition against AK83, the results appeared discordant (Table 1). Indeed, the model well explained the competition phenomenon only for those strains that belong to the training set (R^2^ = 0.99), while it was unreliable for the test set (R^2^ = 0.11). Specifically, AK58 and GR4 performances were underestimated (Fig. 2B). An intermediate amount of “common” k-mers significantly associated with competition phenotype against AK83 was achieved (Table 1).

The modeling analyses were also trained, creating three different random combinations of training and test set. The coefficient of determination R^2^ of the linear regression models of the Rm1021 and AK83 datasets significantly changed compared to the previous analyses, indicating that the models were strongly influenced by the random split of samples into training and test set (Table S3). On the contrary, the coefficients of determination R^2^ of BL225C dataset were less affected by the different combinations of training/test set, indicating that this competition phenotype was well represented by the model for both training and test sets (Table S3). Overall, these results suggest that the linear regression model can well predict the most contrasting competition phenotype in BL225C.

### Putative genetic determinants associated with good competition capabilities against *S. meliloti* BL225C

The linear regression model for the competition against BL225C was the most reliable; therefore, we focused on the set of k-mers related to this dataset. We produced a list of k-mers significantly associated with competition phenotypes covering putative genetic variants. K-mers (p-value < 0.05) were mapped on the annotated genomes of 13 tested strains to retrieve the genetic determinants associated with the phenotype of interest. Among the top k-mers (p-value threshold < 0.0001, Table S4), 51 k-mers (with a p-value equal to 1.31 e^-04^) mapped in genomes of the four strains that showed single nodule occupancy higher than 60% (*S. meliloti* GR4 SM11 KH35c and KH46) in the competition test against BL225C (Table S4, highlighted in bold). Unlike other top k-mers, the coefficients in the linear regression model of these 51 k-mers have all positive values, indicating that there is a positive correlation between the presence of k-mer found in the sample and the phenotypic value attestation. Consequently, based on the above evidence, we focused on these 51 k-mers to define the genetic determinants putatively associated with good competition capabilities against BL225C. These k-mers tagged 103 predicted protein-coding sequences (CDS) (one k-mer can tag multiple genes, Table S5). Among the 103 CDSs, a set of orthologous genes was identified (Table S4). These orthologous genes hits were mostly tracked in *S. meliloti* GR4, KH35c and KH46 genomes (Fig. S4B) and were predominantly located on the symbiosis-required megaplasmids pSymA (ranging from 93.3% to 100%; Fig. S4G, H and I), in particular in a specific region of 26 kb present in the genome of these three strains only (Fig. 3A). On the contrary, 60% of the detected orthologous gene hits in SM11 genomes were located on the chromid pSymB (Fig. S4J).

**Figure 3.**
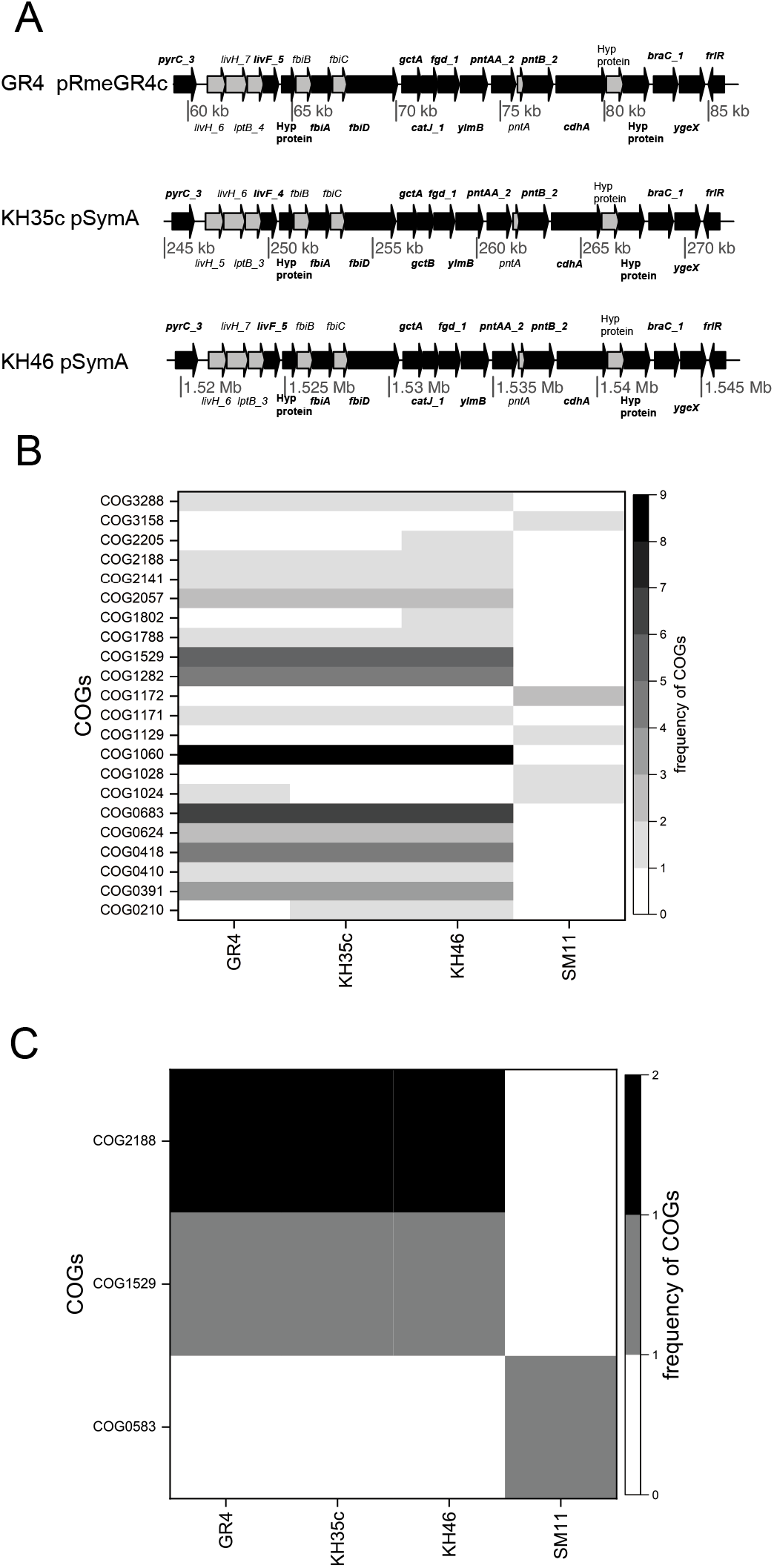
Genetic determinants associated with an increase of competition capabilities. A) k-mers mapping in a region of the symbiotic megaplasmid (pSymA or homologue plasmids) present exclusively in the genome of *S. meliloti* GR4, KH35c and KH46; Genes containing one or more k-mers are indicated with a black arrow, gene annotation is referred to Prokka output. Frequency of candidate functions of B) gene hits and C) regulatory regions identified by 51 best k-mers in the highest competitive strains. The frequency of candidate functions reported as COG annotations (rows) in each strain (columns) is represented by grey-scale shades.

Based on COG annotations, the frequency of orthologous genes cluster hits associated with these k-mers was investigated (Table 2, Table S6). The distribution of candidate function of gene hits within the most competitive strain genomes was not uniform. Enrichment for COG categories E (amino acid transport and metabolism), C (energy production and conversion), H (coenzyme transport and metabolism) and I (lipid transport and metabolism) was found in *S. meliloti* GR4, KH35c and KH46. Notably, the most represented orthologous genes groups are related to coenzyme F_420_ biosynthesis process (COG1060 and COG0391), transmembrane transport via ABC-type systems for branched-chain amino acids (COG0683, COG0410), and pyrimidine nucleotide biosynthetic process (COG0418) (Fig. 3B). Further, a putative caffeine dehydrogenase engaged in pathway of the caffeine transformation via C-8 oxidation (COG1529), and for the two subunits (PntA and PntB) of a presumptive proton-translocating NAD(P) transhydrogenase (COG3288, COG1282) liable for NADPH balancing mechanisms (Table 2) were found. Besides, other presumed functions common to the three strains (GR4, KH35c and KH46) were related to amino-acid degradation/catabolic process (COG1171, COG1788 and COG2057), carbohydrate metabolic process and oxidation-reduction process (COG2141), and transcriptional regulation by GntR-type regulator (COG2188) (Table 2).

**Table 2.** List of functions putatively involved in promoting competing abilities. COG description of gene hits identified by 51 k-mers (p-value 1.13E-04) in the most competitive strains (GR4, KH35c, KH46, SM11). Function/annotation are reported according to the annotation performed with Prokka in this work and/or using original annotation.

| COG ID | COG classes | COG functional category | Prokka annotation / Other annotation | Biological process |
| --- | --- | --- | --- | --- |
| COG0210 | L | Superfamily I DNA or RNA helicase | ATP-dependent DNA helicase PcrA | Mismatch repair, Nucleotide excision repair |
| COG0391 | GH | Archaeal 2-phospho-L-lactate transferase/Bacterial gluconeogenesis factor, CofD/UPF0052 family | Putative phosphoenolpyruvate transferase FbiA | Coenzyme F420 biosynthesis |
| COG0410 | E | ABC-type branched-chain amino acid transport system, ATPase component | High-affinity branched-chain amino acid transport ATP-binding protein LivF | High-affinity branched-chain amino acid transport |
| COG0418 | F | Dihydroorotase | Dihydroorotase PyrC | Pyrimidine nucleotide biosynthesis |
| COG0624 | E | Acetylornithine deacetylase/Succinyl-diaminopimelate desuccinylase or related deacylase | Probable N-formyl-4-amino-5-aminomethyl-2-methylpyrimidine deformylase | Unknown function |
| COG0683 | E | ABC-type branched-chain amino acid transport system, periplasmic component | Leucine-, isoleucine-, valine-, threonine- and alanine-binding protein Brac | Branched-chain amino acid transport |
| COG1024 | I | Enoyl-CoA hydratase/carnithine racemase | Putative fatty acid oxidation complex subunit alpha - enoyl-CoA hydratase FadJ | Fatty acid metabolism |
| COG1028 | IQR | NAD(P)-dependent dehydrogenase, short-chain alcohol dehydrogenase family | Putative NAD-dependent glycerol dehydrogenase | Glycerol metabolic process |
| COG1060 | H | 2-iminoacetate synthase ThiH (tyrosine cleavage enzyme, thiamine biosynthesis) | FbiC FO synthase | Coenzyme F0 biosynthesis |
| COG1129 | G | ABC-type sugar transport system, ATPase component | Ribose import ATP-binding protein RbsA, CUT2 family | Ribose transmembrane transport |
| COG1171 | E | Threonine dehydratase | Diaminopropionate ammonia-lyase | Cellular amino acid catabolic process |
| COG1172 | G | Ribose/xylose/arabinose/galactoside ABC-type transport system, permease component | Autoinducer 2 import system permease protein LsrD | AI-2 transport system |
| COG1282 | C | NAD/NADP transhydrogenase beta subunit | NAD(P) transhydrogenase subunit beta PntB | Oxidation-reduction process<br>Nicotinate and nicotinamide metabolism |
| COG1529 | C | CO or xanthine dehydrogenase, Mo-binding subunit | Putative caffeine dehydrogenase subunit alpha | Oxidation-reduction process |
| COG1788 | I | Acyl CoA:acetate/3-ketoacid CoA transferase, alpha subunit | Glutaconate CoA-transferase GctA subunit A | Glutamate catabolic process (via hydroxyglutarate) |
| COG1802 | K | DNA-binding transcriptional regulator, GntR family | HTH-type transcriptional repressor RspR | DNA-binding transcriptional regulation |
| COG2057 | I | Acyl CoA:acetate/3-ketoacid CoA transferase, beta subunit | Glutaconate CoA-transferase GctB subunit B | Glutamate catabolic process (via hydroxyglutarate) |
| COG2141 | HR | Flavin-dependent oxidoreductase, luciferase family (includes alkanesulfonate monooxygenase SsuD and methylene tetrahydromethanopterin reductase) | F420-dependent glucose-6-phosphate dehydrogenase | Carbohydrate metabolic process, oxidation-reduction process |
| COG2188 | K | DNA-binding transcriptional regulator, GntR family | Putative transcriptional repressor | DNA-binding transcriptional regulation |
| COG2205 | T | K<sup>+</sup>-sensing histidine kinase KdpD | two-component system sensor histidine kinase KdpD | Two-component regulatory system K<sup>+</sup> sensing that regulate kdpABCF operon for Potassium transport |
| COG3158 | P | K<sup>+</sup> transporter | Low affinity potassium transport system protein Kup | Potassium ion transport |
| COG3288 | C | NAD/NADP transhydrogenase alpha subunit | NAD(P) transhydrogenase subunit alpha PntA | Oxidation-reduction process<br>Nicotinate and nicotinamide metabolism |

**Table 3.** List of regulatory regions putatively involved in promoting competing abilities. COG description of supposed target orthologous genes of regulatory region hits identified by 10 k-mers (p-value 1.13E-04) in the most competitive strains (GR4, KH35c, KH46, SM11).

| COG ID | COG classes | COG functional category | Prokka annotation / Product | Biological process |
| --- | --- | --- | --- | --- |
| COG2308 | S | Uncharacterized conserved protein, circularly permuted ATPgrasp superfamily | Uncharacterized putative protein | Function unknown |
| COG2188 | K | DNA-binding transcriptional regulator, GntR family | Putative transcriptional repressor | DNA-binding transcriptional regulation |
| COG0583 | K | DNA-binding transcriptional regulator, LysR family | HTH-type transcriptional regulator DmlR | Transcriptional regulator of dmlA r(aerobic growth on D-malate as the sole carbon source) |
| COG1529 | C | CO or xanthine dehydrogenase, Mo-binding subunit | Putative caffeine dehydrogenase subunit alpha | Oxidation-reduction process |

The number of orthologous genes groups with functional annotation tagged by the 51 best k-mers was lower in SM11. Except for orthologous group related to fatty acid metabolic process (COG1024), the candidate functions of gene hits identified in SM11 were exclusive. Among those putative ABC-type ribose import protein RbsA (COG1129), hypothetical autoinducer 2 import system permease (COG1172), and a putative NAD-dependent glycerol dehydrogenase, belonging to large short-chain dehydrogenases/reductases family (SDR) (COG1028) were identified (Table 2). Among the 103 CDSs tagged by the 51 best k-mers, predicted protein-coding sequences (CDS) with no assigned function were also identified (Tables S5). A large part of these tagged CDS were identified in the SM11 genome and almost entirely located on the SM11 chromosome (Fig. S4A and F). In contrast, CDSs that were found in GR4, KH35c and KH46 genomes were located on homologs of the symbiosis-required megaplasmids pSymA (Fig. S4C, D and E). This last finding suggests the presence of a still remarkable amount of unknown genetic determinants potentially involved in the competition phenotype that require further investigation.

In the mapping procedure of k-mers related to the competition against BL225C, regulatory regions were also analyzed. We considered *bonafide* promoter sequences when hits mapped within 600 nucleotides upstream from CDS start, as previously reported (30). Ten of the 51 k-mers analyzed pinpointed 15 regulatory regions putatively associated with competition phenotype in *S. meliloti* GR4, SM11, KH35c and KH46 (Table S7).

Seven regulatory region hits were associated with CDS with no assigned function (Table S7). These regulatory region hits were tracked exclusively in GR4 and SM11 genomes (Fig. S5A), and were mainly located on the chromosome of the two strains (Fig. S5C and D).

Eight regulatory region hits of putatively orthologous genes target were identified (Table S7, Fig. S5B). These eight regulatory region hits were entirely located on symbiosis-required megaplasmids pSymA in *S. meliloti* GR4, KH35c and KH46 (Fig S5E, F and G) and on pSymB of *S. meliloti* SM11 (Fig S5H).

In GR4, KH35c and KH46, the regulatory region hits are associated with genes encoding for proteins whose functions (COG1529 and COG2188) was previously observed (Fig. 3B and C), suggesting a role for genes in these COGs in determining competition phenotypes. In SM11, the regulatory regions hits are associated with a LysR-type orthologous gene (transcriptional regulation) and with a gene belonging to COG2308 possibly involved in the biosynthesis of small peptides.

## DISCUSSION

Rhizobia-legume symbiosis is a paradigmatic example of bacteria-plant association, where bacteria behave as facultative symbionts, colonizing the plant tissue in nodules and establishing intracellular infection. The ability to colonize plant tissue is under selective pressure and rhizobial strains, which efficiently colonize host plants, have been shown to more effectively promote plant growth, giving rise to a partial “fitness alignment” between the host and the symbiont (33, 34). However, in nature, several strains compete for forming symbiotic associations with the host plant and often nodules are simultaneously colonized by different strains, which in turn may have different efficiency in promoting plant growth, some of them behaving as cheaters also (7, 8, 16, 35). Then, measuring the competitiveness for symbiosis and nodule colonization and predicting this phenotype from rhizobial genome sequencing is of paramount importance for understanding the evolution of rhizobia-plant symbiosis and developing effective inoculant strains for increasing agricultural yield of legume crops (8).

Here we addressed the possibility to develop a predictive model of strain competitiveness in the host plant–rhizobial symbiont systems *M. sativa–S. meliloti*. Direct measurements of competitiveness were obtained through the analysis of nodule occupancy. This experimental design pointed out a wide variety of strain response to the diverse competitive conditions, identifying three different competition patterns and outlining a highly complex phenotype that strongly depends on the engaged competitor. Strains used in this work were originally isolated from different *Medicago* species, however they displayed good competition capabilities in nodulation of *M. sativa*, indicating that nodulation competitiveness is not strictly bound to the host genotype. Moreover, an abundant presence of mixed nodules was observed, confirming previous results (7, 11, 36). Mixed nodules were found in all the three sets of experiments suggesting that the possibility to co-infect nodules by different strains could be more widespread than expected. Strains M270 and T073, which were characterized by low nitrogen fixation rates, also showed low competition capabilities. Conversely, the differential response of strains with medium-high nitrogen-fixation efficiency advance the idea that a greater competitive ability is not correlated with high nitrogen fixation efficiency in *M. sativa–S. meliloti* interaction as previously suggested (37, 38). Concerning the competition against Bl225C, except for GR4 and SM11, this assumption seems to be particularly true for highly efficient N-fixer strains (USDA1157, CCMM-B554, RU11/001, 2011, and AK58), which turned out to be medium-weak competitors. Co-persistence strategies in mixed infections based on the exploitation of benefits by less mutualistic strains were previously hypothesized (39). Indeed, for these strains, low competitive abilities coupled with a high percentage of mixed nodules hinted at the possibility that exploitation of more effective nitrogen fixer strains in mixed nodules by competitor may occur.

For the evaluation of the genetic determinants responsible of an increased competition phenotype, we used a method based on the use of k-mers, in GWAS, this kind of approaches are progressively increased in recent years (18, 40, 41), thanks to their flexibility in capturing different types of genetic variants and overcoming the alignment of sequence to a reference genome. PhenotypeSeeker is one of the up-to-date tools that utilize machine learning for predicting phenotypes from the sole genomes sequences (32). This software combines the possibility to identify k-mers as signatures of competition phenotype with the training of a predictive linear regression model. Besides supplying statistical information that establishes the association degree between the presence of the k-mers in samples with their assessed competition phenotype, the evaluation of predicting power of built models can also be a hint of the degree of the association of top k-mers with phenotype. The results of the analyses with different combinations of training/test set ensure the robustness of the linear regression model. The influence exerted by different combinations of training/test set on the accuracy of the model, marked that competition phenotypes characterized by either positive or negative trends are less suitable for the regression analysis. In the case of Rm1021 dataset, the negative value of averaged R^2^ indicates that the mean of the phenotype values of training samples have more predictive power on the test set than the model itself. In contrast, the heterogeneous phenotypes ascertained in competition against BL225C might be well predicted by the trained model.

The evaluation of the predictive power of k-mers involved in model training is also a clue of the degree of the association of the latter to the phenotype competitiveness. Beyond the statistical analyses, the 51 k-mers related to the competition against BL225C that were taken into account are also highly predictive k-mers. Aside from having a significant impact on the outcomes of the machine-learning model, these k-mers, which featured a positive value of coefficients in the regression model, are positively correlated with the attested phenotype. Moreover, since they mapped in genomes of the four most competitive strains, these 51 k-mers can be considered as the most informative k-mers, allowing us to tag the genetic variants associated with remarkable competition capabilities.

The largest part of genes putatively associated with competitiveness was harboured by the megaplasmid pSymA (or pSymA homologs depending on the strain). Megaplasmid pSymA is the genomic element carrying all the genes necessary for symbiosis (e.g. *nod*, *fix* and *nif* genes) (42). Moreover, it was shown to harbour the largest part of genomic diversity in *S. meliloti* (43), then possibly largely contributing to the phenotypic diversity among strains and, on the basis of the obtained results, it could be linked to competition capabilities too.

Previous studies highlighted the importance of exopolysaccharide production, motility and signalling for symbiosis establishment and competition (25, 27). The genes (with functional annotation) found in this work, putatively associated with competitiveness, are mostly related to biosynthesis and transport functions. Many k-mers are related to COG1060 that, together with COG0391, is linked to the presence of the *fbi* operon in KH46, KH35c and GR4 strains only. The *fbi* operon is widely distributed in aerobic soil bacteria and it is responsible for the synthesis of the functional versatile redox factor F_420_ (44, 45). This cofactor is involved in the redox modification of many organic compounds facilitating low-potential two-electron redox reactions (44, 45). The presence of this cofactor is linked to several important processes such as persistence, antibiotic biosynthesis (tetracyclines, lincosamides and thiopeptides) and pro-drug activation (46); all functions that could play a central role in the competitiveness.

Another group of COGs highly represented in our dataset is related to ABC transporters. It is well known that *S. meliloti* genome encodes for a large number of ABC uptake and export systems (47, 48). This feature is probably linked to selective adaptation to oligotrophic soils and rhizospheric conditions (48, 49). According to our association analysis, a group of k-mers tagged putative genes encoding for ATPase (COG0410) and permease (COG0683) subunits of branched-chain amino acids ABC transport complex Bra/Liv (50). In *S. meliloti*, a double mutant for the two main amino acid ABC transport complexes (*aap bra*) showed no reduction in N_2_-fixation efficiency as well as no influence on the plant phenotype, suggesting the idea that in bacteroids a branched-chain amino acids auxotrophy, called “symbiotic auxotrophy”, does not occur (51). However, an attenuated competitive phenotype was found in *S. meliloti* mutated in *livM* gene, which encodes for the permease subunit of Bra/Liv complex (52). It is then reasonable to suppose that this complex may provide a noteworthy benefit in the competition dynamics ensuring a higher supply of amino acids in free-living rhizospheric conditions and increasing strain competitiveness (52, 53). Other COGs related to ABC transporter are COG1129 and COG1172. Proteins grouping in COG1129 are ATPase component of an ABC-type ribose import system. In *Rhizobium leguminosarum*, a putative ribose ABC transporter (RbsA, RL2720) is induced by the presence of arabinogalactan, and it is specifically overexpressed in alfalfa rhizosphere (49). COG1172 contains ribose/xylose/arabinose/galactoside ABC-type transport system permease component remarking the importance of efficient carbon uptake in the rhizosphere to outcompete other bacteria. COG1172 is also related to the import of autoinducers signalling molecules in the quorum sensing process, whose connection with *S. meliloti* competitive behavior is well known (54).

Several COGs are related to the metabolism of different compounds: pyrimidine (COG0418; dihydroorotase), glutamate (COG1788 and COG2057; glutaconate CoA transferase), amino acids (COG1171; threonine dehydratase), fatty acids (COG1024; enoyl-CoA hydratase/carnitine racemase) and glycerol (COG1028; glycerol dehydrogenase), reinforcing the importance of metabolic versatility to increase adaptation to the rhizosphere. Indeed, in *R. leguminosarum* bv. *viciae* a deficiency for glycerol transport or utilization leads to reduced competitiveness for nodulation (55).

At last, a group of k-mers fell within transcriptional regulation genes (COG2188, COG1802 and COG0583) suggesting their possible involvement for a fine-tuning bacterial response to the presence of other competitors and/or for a quick response to variation of the external conditions. Other COGs retrieved with our approach have less clear connections with competitiveness and will require further studies to infer their possible role in this process: COG1282/3288 (NAD(P) transhydrogenase) that may be involved in removing reactive oxygen species, COG0210 (helicase) and COG1529 (caffeine dehydrogenase).

A substantial part of k-mers maps on hypothetical genes with unknown functions, indeed many genes required for rhizobial adaptation to the rhizosphere are not characterized yet. Transcriptomic analysis of rhizobia isolated from the rhizosphere showed the expression of many hypothetical genes (49). This finding suggests that there is still to discover in the pangenome of *S. meliloti* a number of functions potentially important in the fitness associated with the symbiotic interaction and possibly in plant growth promotion.

Rhizobial competitiveness is a cornerstone for plant colonization; therefore, the selection of highly competitive/efficient rhizobia is fundamental for sustainable agricultural production. Here, we have reported on the feasibility and reliability of using a k-mer-based GWAS approach to detect genes associated with this complex quantitative phenotype in the plant symbiont *S. meliloti*. Several functions contribute to ameliorate competitiveness indicating that many different bricks, increasing rhizobial versatility, pave the way for success in competition. This approach may provide the basis for large-scale screening of putative competitiveness capabilities among pairs of strains on the basis of genome sequences. Interestingly, the evidence that most of the genes putatively associated with competition resides on the megaplasmid pSymA can offer the possibility to extend the creation of *ad hoc* hybrid strains by mobilizing the pSymA megaplasmid from different hosts (56) in the aim to develop novel ameliorated inoculants (8).

## MATERIAL AND METHODS

### Bacterial strains, plasmids and growth conditions

The strains and plasmids used in this work are listed in Table S7. *Escherichia coli* strains were grown in liquid or solid Luria Bertani (LB) medium (Sigma Aldrich) at 37°C (57), supplemented with tetracycline (10 μg/ml). *Sinorhizobium meliloti* strains were cultured in broth or agar tryptone yeast (TY) medium with 0.2 g/l CaCO_3_ at 30° C (58), supplemented with streptomycin (500 μg/ml in broth and agar media), rifampicin (50 μg/ml) and tetracycline (1 μg/ml in liquid broth medium, 2 μg/ml in agar medium), when necessary.

### Competition assay

*Medicago sativa* (cv. Maravigliosa) plantlets were germinated and grown as described in nodulation assays section (Supplementary methods). *S. meliloti* strains were grown at 30° C to late exponential phase (OD_600_ = 0.6 - 0.8) in TY with opportune antibiotics; then each culture was washed 2 times in Nitrogen-free solution and diluted to a final concentration of 5×10^4^ CFU/ml. The inocula mixtures were prepared with equal volumes of cellular suspensions of two different fluorescent-tagged strains. A total of 39 competition experiments were set up (13 GFP-tagged strains x 3 RFP-tagged strains). Six plants for each competition experiment were inoculated with 1 ml of inocula mixtures *per* seedling. After 4 weeks, nodule fluorescence was detected using a fluorescence stereomicroscope, Stereo Discovery V12 (Zeiss; Germany, Oberkochen), equipped with a CCD camera controlled by Axiovision software for image acquisition. All fresh nodules of each plant were individually exposed with filters for GFP (Zeiss Filter Set 38HE, excitation 470/40 emission 525/50) and DsRed (Zeiss Filter Set 43HE, excitation 550/25 and emission 605/70). The images obtained were processed with ImageJ software (59). The number of green nodules (occupied/produced by GFP-tagged strains), red nodules (occupied/produced by RFP-tagged strains) and mixed nodules (occupied/produced by both fluorescent-tagged strains) were counted for each plant of each competition experiment. The nodule occupancy was expressed as the ratio of the number of nodules (green, red or mixed) by the total number of nodules present on the roots of each plant. Statistical analyses of data were performed with nonparametric Kruskal-Wallis and Dunn test post-hoc by using *FSA* and *rcompanion* packages of Rstudio software (60).

### PhenotypeSeeker analysis

The single nodule occupancy of the strains, assessed in the three competition experiments, were converted in a continuous matrix of equivalent values between 0 and 1 for each dataset. The FASTA genome sequences of 13 strains and obtained matrices were used as input to count all k-mers for each set of competition. The k-mer length was set to 13 nucleotides. In the first filtering step, the k-mers that were present in or missing from less than two samples (“— min 2—max 2”; default) were rejected. The clonal population structure correction was performed. The k-mers were tested for the analyses of association with the phenotype according to the weighted Welch two-sample t-test, and the k-mers with a p-value higher than 0.05 were automatically discarded. The linear regression models were achieved using only top lowest p-valued k-mers for all three datasets, with default parameters. For the first regression analyses, the strains were split into a training test (KH46, CCMM-B554, T073, Rm41, HM006, 2011, USDA1157, SM11 and RU11/001) and test set (AK58, KH35c, GR4, M270). In the second regression analyses, three different random combinations of training and test set were used, and the model evaluation metrics were averaged over 3-fold train/test splits. The 3-fold explicitly indicates that each strain was once included into test set and twice included into training set.

## SUPPLEMENTAL MATERIAL

**Supplementary methods**

**Fig. S1**

**Fig. S2**

**Fig. S3**

**Fig. S4**

**Fig. S5**

**Table S1**

**Table S2**

**Table S3**

**Table S4**

**Table S5**

**Table S6**

**Table S7**

**Table S8**

## ACKNOWLEDGEMENTS

Erki Aun and Maido Remm were funded by the institutional grant IUT34-11 from the Estonian Ministry of Education and Research and the EU ERDF grant No. 2014-2020.4.01.15-0012 (Estonian Center of Excellence in Genomics and Translational Medicine).

A.B., F.P., A.M. and C.V. conceived and planned the research; A.B. and F.P. performed competition assays and symbiotic assay; E.Az. and F.D. contributed to competition assays; E.Au. and M.R performed PhenotypeSeeker analysis and contributed to linear regression model interpretation; A.B. and G.B. performed the bioinformatics analyses; A.B., F.P., A.M, L.G., and C.V. interpreted data; A.B., F.P., A.M. prepared the manuscript; all authors participated in editing the manuscript.

We declare no competing interests.

