## Supplemental material for "Competitiveness prediction for nodule colonization in *Sinorhizobium meliloti* through combined *in vitro* tagged strain characterization and genome-wide association analysis"

SUPPLEMENTARY MATERIAL

**Supplementary methods**

**- Construction of *S. meliloti* fluorescently tagged strains**.

**- Nodulation and acetylene reduction assays**

**- Annotation and phylogenetic analyses.**

**- Mapping procedure.**

**Supplementary figures**

**Figure S1. Evolutionary relationships and pangenome of strains used as competitors**. A) The evolutionary history was inferred using the UPGMA method on core genes concatenamer alignment. The optimal tree with the sum of branch length = 0.02977589 is shown. The percentage of replicate trees in which the associated taxa clustered together in the bootstrap test (10000 replicates) is shown next to the branches. The evolutionary distances were computed using the Maximum Composite Likelihood method and are in the units of the number of base substitutions per site. There were a total of 3998704 positions in the final dataset. B) Heatmap showing gene presence (dark blue) or absence light blue in each strain, the tree on the left was build on the basis of presence/absence of genes. C) Histogram showing the frequency of genes depending on the number of genomes. D) Pie chart displaying all the genes present in the pangenome and their breakdown in the different genomes.

**Figure S2. Nodulation assay and nitrogen fixation efficiency with single strains**. A) Number of nodules/plant; B) Epicotyl length; C) Plant dry weight; D) Acetylene reduction assay (ARA). Different letters indicate significant differences between treatments ($p < 0.05$).

**Figure S3. Number of nodules in the three competitions.** A) Total nodules *per* plant. B) Total mixed nodules *per* plant. Different letters indicate significant differences between treatments ($p < 0.05$).

**Figure S4. Distribution of 51 best k-mers tagged-CDSs in S. *meliloti* GR4, KH35c, KH46 and SM11.** A) Distribution of unannotated CDSs among four *S. meliloti* strains and distribution of unannotated CDSs among replicons of *S. meliloti* strains C) GR4, D) KH35c, E) KH46 and F) SM11. B) Distribution of orthologous genes hits among four strains *S. meliloti* strains and distribution of orthologous genes among replicons of *S. meliloti* strains G) GR4, H) KH35c, I) KH46 and J) SM11.

**Figure S5. Distribution of 10 k-mers tagged-putative regulatory regions in *S. meliloti* GR4, KH35c, KH46 and SM11.** A) Distribution of regulatory region hits of unannotated CDSs among four *S. meliloti* strains and among replicons of *S. meliloti* strains C) GR4 and D) SM11. B) Distribution of putative regulatory region hits of orthologous gene hits among four *S. meliloti* strains and among replicons of *S. meliloti* strains E) GR4, F) KH35c, G) KH46 and H) SM11.

**Supplementary tables**

**Table S1**. *Sinorhizobium meliloti* strains used in this work.

**Table S2.** Single nodule occupancy of *S. meliloti* tested strains in competition experiments versus S. meliloti strains Rm1021, AK83 and BL225C. Different letters indicate statistically significant differences (Kuskal-Wallis and Dunn test, $p<0.05$) within a competition assay (columns; vs BL225C, vs AK83, vs Rm1021).

**Table S3**. Linear regression models for the three competition experiments, performed by PhenotypeSeeker with 3-fold train/test splits of samples. The averaged model evaluation metrics of both training and test set are reported.

**Table S4.** List of top k-mers (raw data k-mers).

**Table S5.** Genes hits identified by 51 best k-mers (raw data kmers - gene position).

**Table S6.** List of COGs codes.

**Table S7.** Regulatory region hits identified by 10 best k-mers.

**Table S8**. Strains and plasmids used in this work

**Supplementary methods**

**Construction of *S. meliloti* fluorescently tagged strains**. *S. meliloti* strains were tagged with green fluorescent protein (GFP) or red fluorescent protein (RFP). Donor *E. coli* S17-1 strains containing plasmids pHC60 (harboring a constitutively expressed GFP; (1)) or pBHR mRFP (harboring a constitutively expressed RFP; (2)) were used for biparental conjugations with rifampicin-resistant derivatives *S. meliloti* strains. Spontaneous rifampicin derivative *S. meliloti* strains were isolated by plating aliquots of 100 µl of cell suspension of $10^9$ cells on agar TY medium with rifampicin (50 µg/ml). Conjugal transfer was performed as previously described (3)

**Nodulation and acetylene reduction assays**. *Medicago sativa* (cv. Maravigliosa) seedlings were surface sterilized with 70% ethanol for 1 min, rinsed with sterile dd$H_2O$, treated with 2.5% sodium hypochlorite for 5 min and washed 20 times with sterile dd$H_2O$. Sterilized seeds were then let germinate on the cover of sterile plastic Petri dishes upside down for 4 days in the dark at room temperature. Seedlings were transferred in plastic pots containing a sterilized mixture of sand and vermiculite (ratio 2:3) and supplied with 120 ml of sterilized Nitrogen-free solution (1mM $CaCl_2$ $2H_20$, 0.1 mM KCl, 0.8 mM $MgSO_4$ $7H_2O$, 10 µM Fe EDTA, 35 µM $H_3BO_3$, 9 µM $MnCl_2$ $4H_2O$, 0.8 µM $ZnCl_2$, 0.5 µM $Na_2MoO_4$ $2H_2O$, 0.3 µM $CuSO_4$ $5H_2O$, 3.68 mM $KH_2PO_4$, 4 mM $Na_2HPO_4$ pH=6.5) (4). Seedlings were grown for 3 additional days before inoculation with *S. meliloti* strains. The strains were grown at 30 °C to late exponential phase ($OD_{600}$ < 0.6 - 0.8), washed 2 times in Nitrogen-free solution and then adjusted to an $OD_{600}$ = 0.05 in Nitrogen-free solution. Nine plants for strains were inoculated with aliquots of 500 µl of cell suspension of $5\times10^7$ CFU/ml, and grown in a growth chamber maintained at 23 °C with a 16-h photoperiod. The same amount of Nitrogen-free solution was added to negative control plants (C-). After 28 days, the epicotile length, number of nodules and dry weight were measured.

For the acetylene-reduction assay, *M. sativa* plants were grown as described above. After 28 days, plants were collected in 100 ml glass flasks (3 plants/flask) and sealed with gas-thigh silicone caps. Aliquots of 10 ml of acetylene were injected into the flasks and, after 40 min, the ethylene concentration was measured by using a 7890B gas chromatograph system (Agilent technologies; California, USA), equipped with a 5975 Mass selective detector. Chromatographic analyses were performed in the following conditions: initial temperature, 40°C (isocratic for 10 min), gas flow (helium) 4 ml/min, injection 500 µl (gas syringe) at a split ratio of 5:1. Nitrogen fixation rates were expressed in nanomoles of produced ethylene *per* hour, *per* plant.

Statistical analysis of data was performed with Rstudio software (5). Shapiro test was performed to evaluate data distribution; ANOVA and Tukey post-hoc test or nonparametric Kruskal-Wallis and Dunn test post-hoc were performed using *FSA* and *rcompanion* packages.

**Annotation and phylogenetic analyses.** *S. meliloti* genomes were retrieved from the NCBI Genome Database (GenBank codes are reported in Table S1). Genome annotation of 13 *S. meliloti* strains was completed using Prokka (version 1.13) bacterial genome annotation tool (6).The pangenome of the 13 *S. meliloti* strains was constructed with Roary 3.11.3 (7) using default settings to construct a whole-genome phylogeny. Core genes alignment, obtained with Roary, was used to infer the evolutionary relationship of the strains tested as competitors. The evolutionary distances were computed using the Maximum Composite Likelihood method and are in the units of the number of base substitutions per site (8). The evolutionary history was reconstructed using the UPGMA method (bootstrap test of 1000 replicates). All ambiguous positions were removed for each sequence pair (pairwise deletion option). All evolutionary analyses were conducted using MEGA X software (9).

**Mapping procedure.** For each competing strain tested, the genome position of k-mers associated with the phenotype (competition against BL225C strain) was detected using the R package Biostrings (version

2.54) (10). Only k-mers aligning without mismatches or gaps on the positive or negative strand of the reference were taken into account to reflect the pipeline used by PhenotypeSeeker, which does not allow for mismatches. Absolute positions of k-mers were then transformed into relative ones based on genomic annotations following four rules:

1. If a k-mer was mapped inside a gene, its position was set to 0 independently from the strand
2. If a k-mer was mapped outside a gene on the positive strand, its position was adjusted by subtracting the starting position of the nearest gene. Since the starting position of the nearest gene on the plus strand is always greater than the starting position of the k-mer, the relative position will always be a negative value representing the number of bases ahead of the sequence of the gene on the reference genome.
3. If a k-mer was mapped outside a gene on the negative strand, its position was calculated by subtracting the starting position of the k-mer to the ending position of the gene. Analogously to the previous calculation, the starting position of the k-mer will always be greater than the ending position of the gene on the minus strand, thus the relative position of the k-mer will always be a negative value representing the number of bases behind the sequence of the gene on the reference genome.
4. If a k-mer was mapped ahead of a gene on the positive strand and behind a gene on the negative strand (namely "between" two genes oriented in different directions), its position was calculated as reported in 1 and 2. Since both relative positions may be valid they were both reported and considered in downstream analyses.

Relative position obtained were then used to extract the predicted protein-coding sequences (CDS) and regulatory regions mapped by 51 k-mers (with a p-value =1.31 e-04) by selecting those with a relative position equal to 0 and higher than -600 respectively.

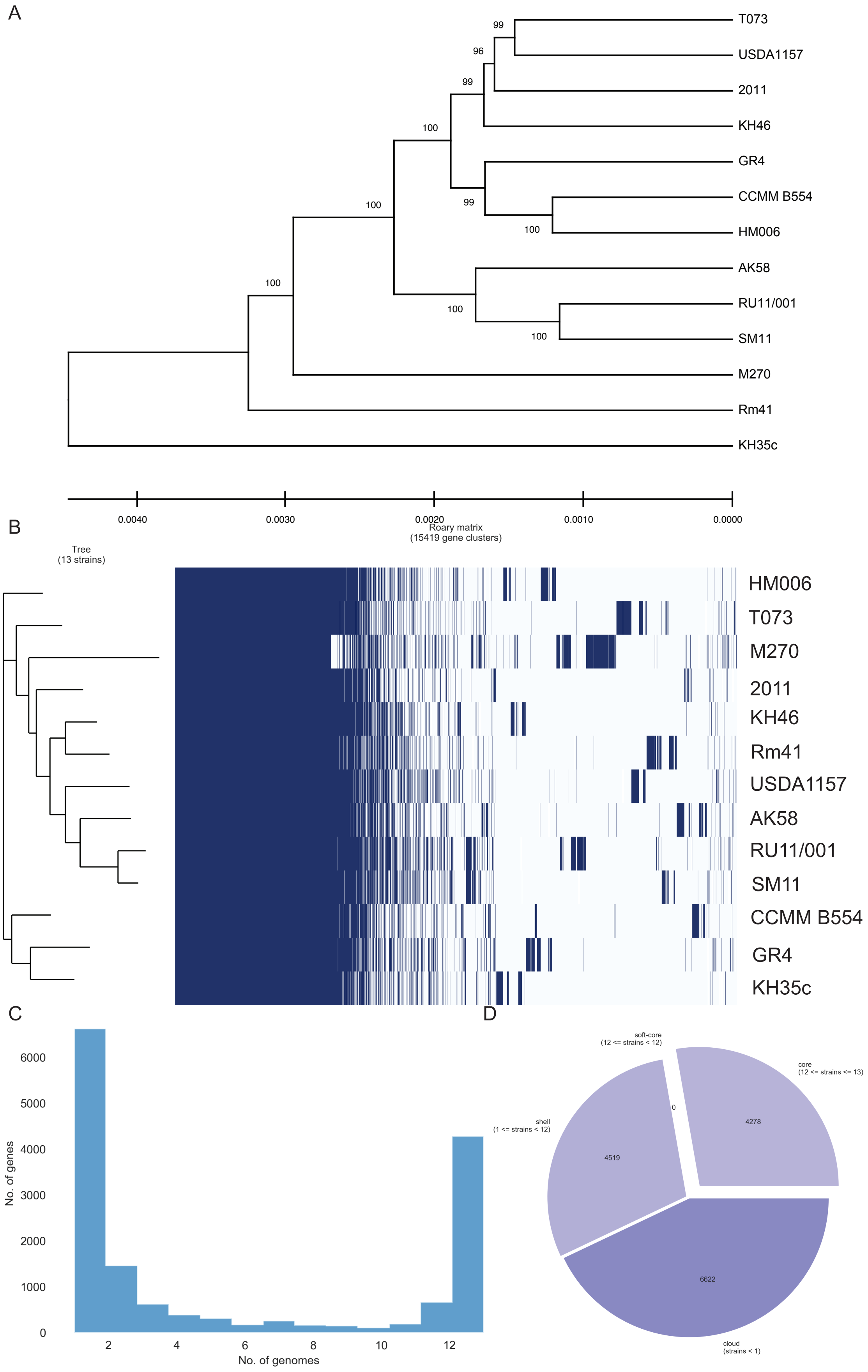
A
T073
USDA1157
2011
KH46
GR4
CCMM B554
HM006
AK58
RU11/001
SM11
M270
Rm41
KH35c
99
96
99
100
100
99
100
100
100
100
0.0040
0.0030
0.0020
0.0010
0.0000
B
Tree
(13 strains)
Roary matrix
(15419 gene clusters)
HM006
T073
M270
2011
KH46
Rm41
USDA1157
AK58
RU11/001
SM11
CCMM B554
GR4
KH35c
C
No. of genes
0
1000
2000
3000
4000
5000
6000
2
4
6
8
10
12
No. of genomes
D
soft-core
(12 <= strains < 12)
core
(12 <= strains <= 13)
shell
(1 <= strains < 12)
cloud
(strains < 1)
0
4278
4519
6622

**Figure S1. Evolutionary relationships and pangenome of strains used as competitors**. a) The evolutionary distances were computed using the Maximum Composite Likelihood method and are in the units of the number of base substitutions per site. The evolutionary history was inferred using the UPGMA method. The optimal tree with the sum of branch length = 0.03043492 is shown. The percentage of replicate trees in which the associated taxa clustered together in the bootstrap test (1000 replicates) are shown next to the branches. There were a total of 4134474 positions in the final dataset. b) heatmap showing gene presence (dark blue) or absence light blue in each strain, the tree on the left was build on the basis of presence/absence of genes. c) histogram showing the frequency of genes depending on the number of genomes. d) pie chart displaying all the genes present in the pangenome and their breakdown in the different genomes.

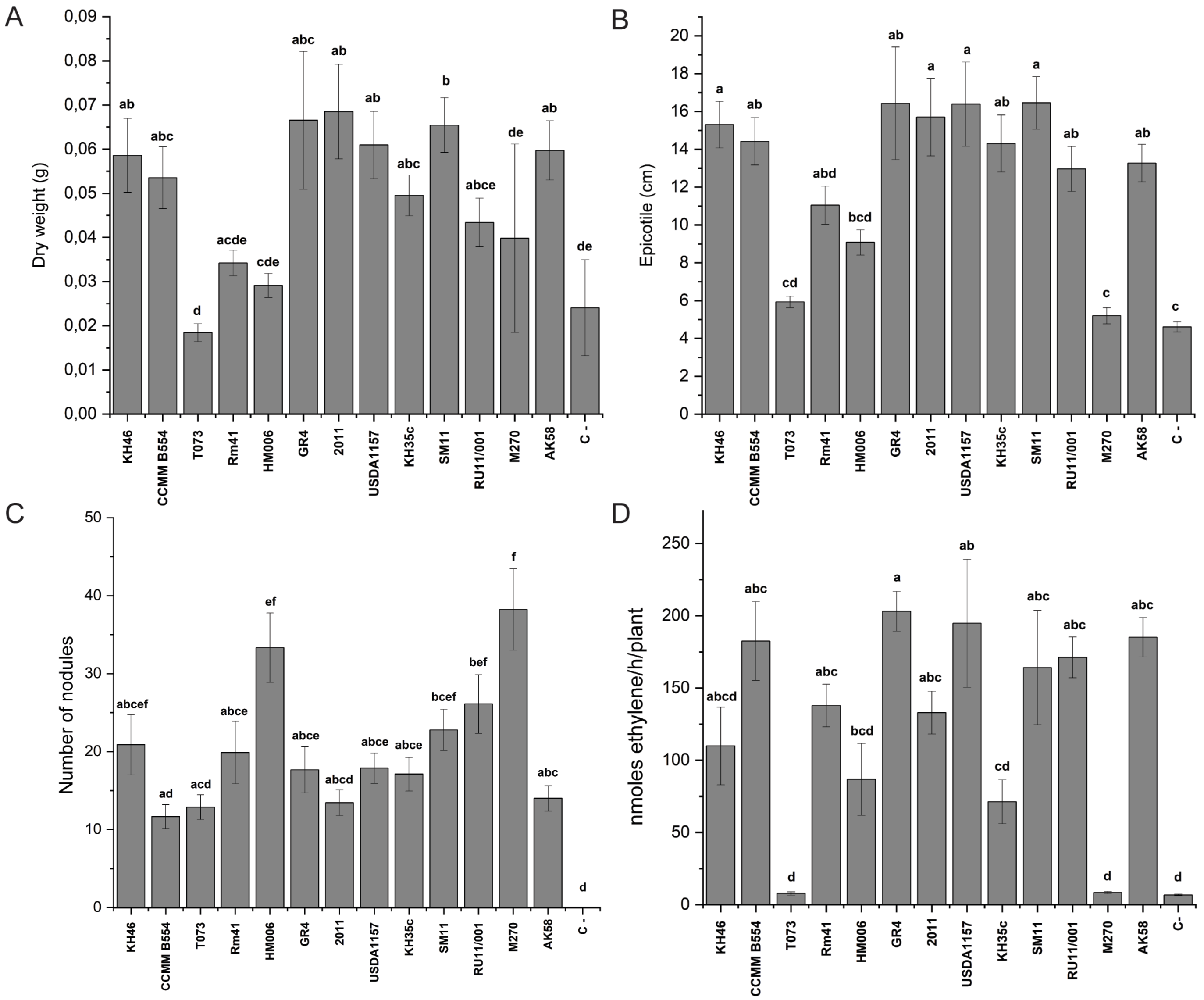


**Figure S2. Nodulation assay and nitrogen fixation efficiency with single strains**. a) Number of nodules/plant; b) Epicotyl length; c) Plant dry weight; d) Acetylene reduction assay (ARA).

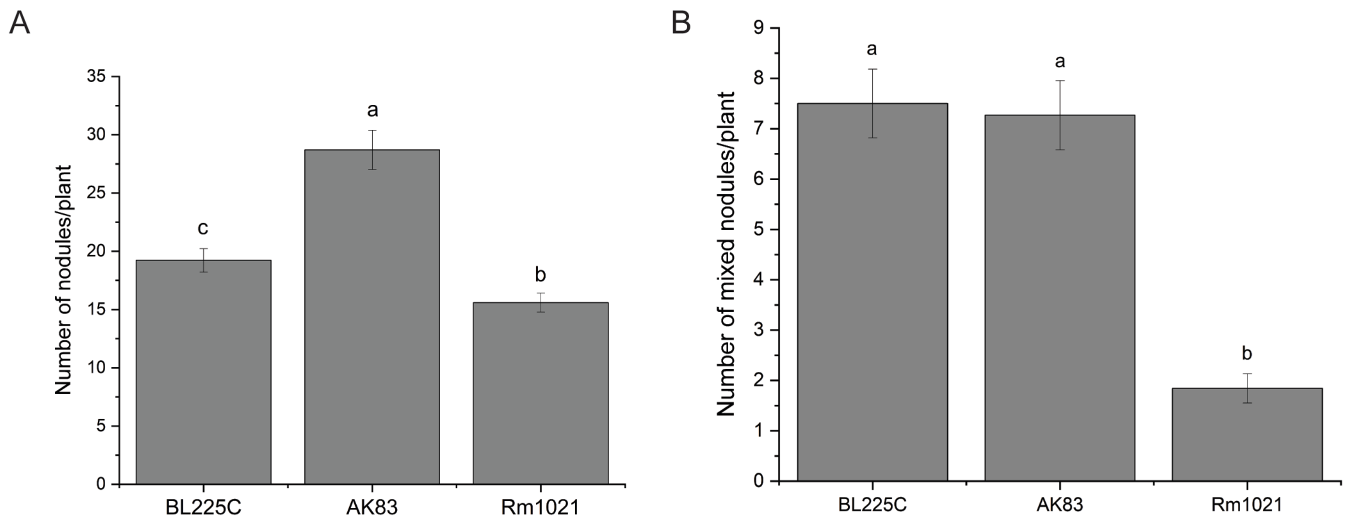


**Figure S3. Number of nodules in the three competitions.** A) Total nodules *per* plant. B) Total mixed nodules *per* plant. Different letters indicate significant differences between treatments ($p < 0.05$).

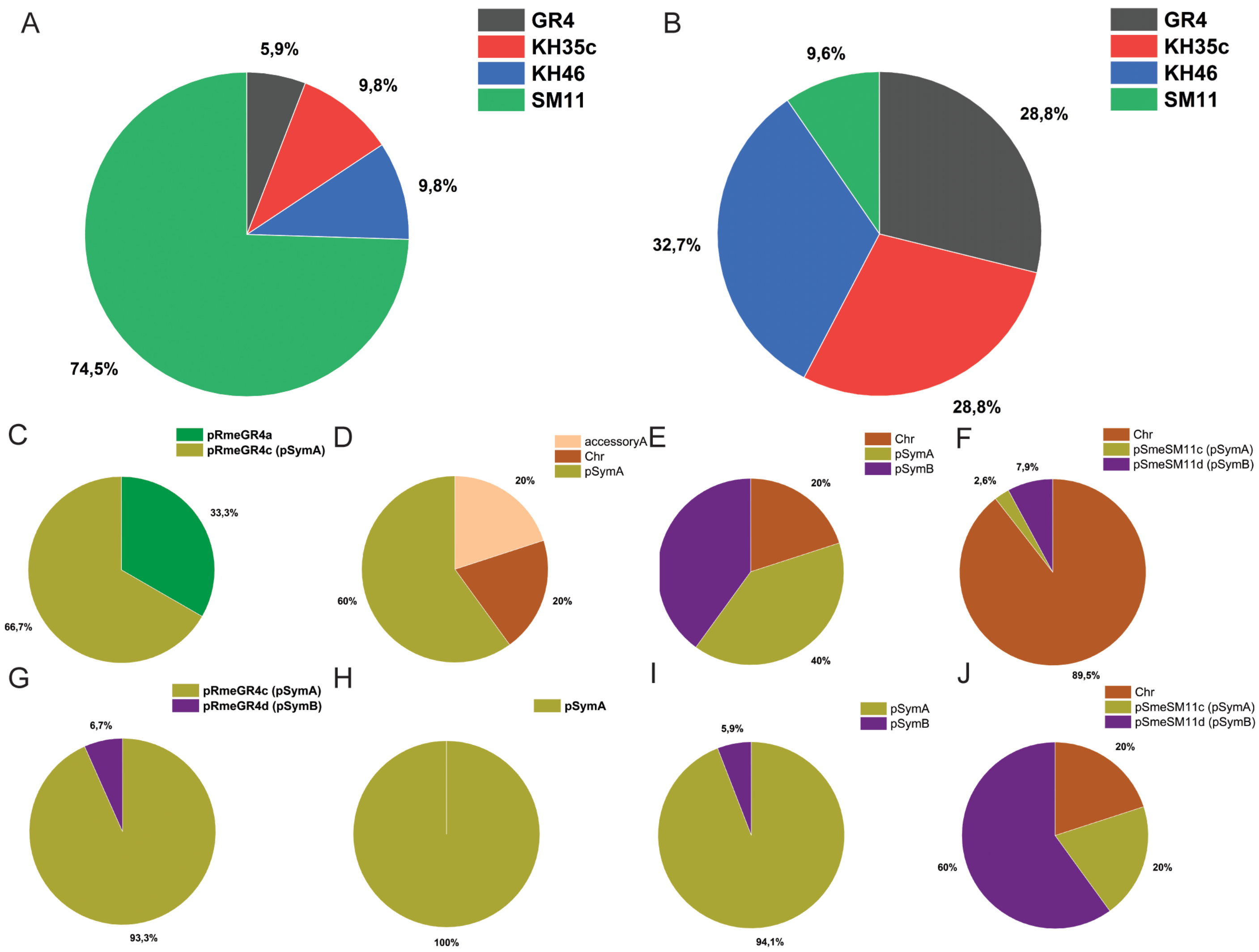


**Figure S4. Distribution of 51 best k-mers tagged-CDSs in *S. meliloti* GR4, KH35c, KH46 and SM11.** A) Distribution of unannotated CDSs among four *S. meliloti* strains and distribution of unannotated CDSs among replicons of *S. meliloti* strains C) GR4, D) KH35c, E) KH46 and F) SM11. B) Distribution of orthologous genes hits among four strains *S. meliloti* strains and distribution of orthologous genes among replicons of *S. meliloti* strains G) GR4, H) KH35c, I) KH46 and J) SM11.

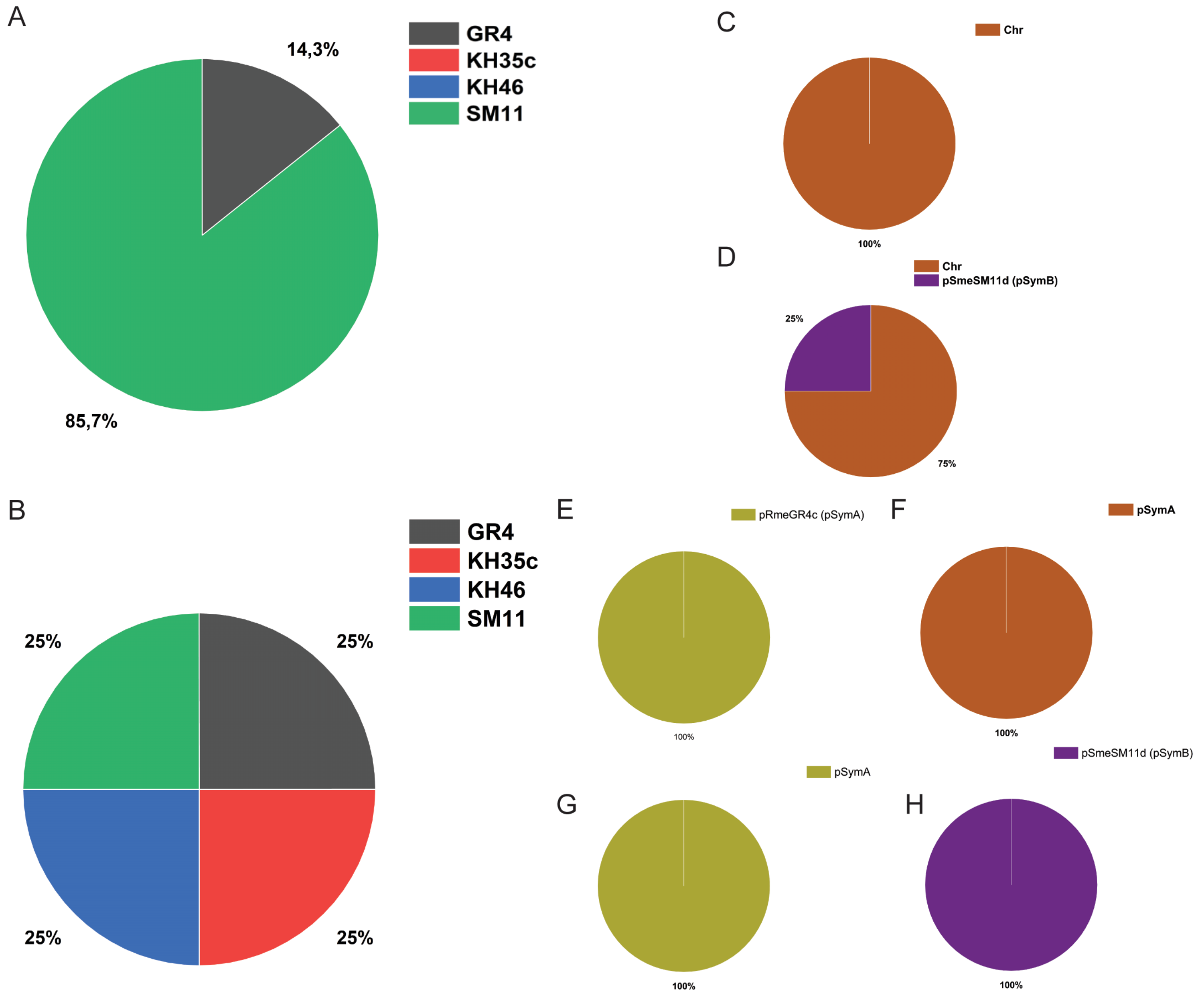


**Figure S5. Distribution of 10 k-mers tagged-putative regulatory regions in *S. meliloti* GR4, KH35c, KH46 and SM11.** A) Distribution of regulatory region hits of unannotated CDSs among four *S. meliloti* strains and among replicons of *S. meliloti* strains C) GR4 and D) SM11. B) Distribution of putative regulatory region hits of orthologous gene hits among four *S. meliloti* strains and among replicons of *S. meliloti* strains E) GR4, F) KH35c, G) KH46 and H) SM11.

**Table S1**. *Sinorhizobium meliloti* strains used in this work.

| Strains | Source/Description | Genbank assembly codes | Reference |
|---|---|---|---|
| AK83 | Geographic location: Kazakhstan; Host: *Medicago falcata* | GCA_000147795.3 (11) | (12) |
| 1021 | SU47 *str*-21 | GCA_000006965.1 (13) | (14) |
| BL225C | Geographic location: Italy; Host: *Medicago sativa* | GCA_000147775.3 (11) | (15) |
| KH46 | Geographic location: France; Host: *Medicago truncatula* | GCF_002197465.1 (16, 17) | (16) |
| CCMM B554 | Geographic location: Morocco; Host: *Medicago arborea* | GCA_002215195.1 (18) | (19) |
| T073 | Geographic location: Tunisia; Host: *Medicago truncatula* | GCA_002197145.1 (16, 17) | (16) |
| Rm41 | Geographic location: Hungary; Host: *Melilotus/Medicago* | GCA_000304415.1 (20) | (21) |
| HM006 | Geographic location: France; Host: *Medicago truncatula* | GCA_002197165.1 (16, 17) | (16) |
| GR4 | Geographic location: Spain ; Host: agricultural field | GCA_000320385.2 (22) | (22) |
| 2011 | SU47 | GCA_000346065.1 (23) | - |
| USDA1157 | Geographic location: USA, California; Host: *Medicago sativa* | GCF_002197025.1 (17) | (17) |
| KH35c | Geographic location: France; Host: *Medicago truncatula* | GCA_002197105.1 (16, 17) | (16) |
| SM11 | Geographic location: Germany; Host: agricultural field | GCA_000218265.1 (24) | (25) |
| RU11/001 | Geographic location: Germany; Host: *Medicago sativa* | GCA_001050915.2 (26) | (27) |
| M270 | Geographic location: Jordan; Host: *Medicago truncatula* | GCA_002197085.1 (16) | (16) |

| | | | |
|---|---|---|---|
| AK58 | Geographic location: Kazakhstan; Host: *Medicago falcata* | GCA_000473425.1 (28) | (12) |

**Table S2.** Single nodule occupancy of *S. meliloti* tested strains in competition experiments versus S. meliloti strains Rm1021, AK83 and BL225C. Different letters indicate statistically significant differences (Kuskal-Wallis and Dunn test, $p<0.05$) within a competition assay (columns; vs BL225C, vs AK83, vs Rm1021).

| **Strains** | **vs BL225C** | **vs AK83** | **vs Rm1021** |
|---|---|---|---|
| AK58 | 41.2% $^{abcd}$ | 45.7% $^{a}$ | 82.7 % $^{cd}$ |
| CCMM B554 | 42.9% $^{abc}$ | 28.6% $^{ab}$ | 65.0 % $^{acd}$ |
| GR4 | 66.7% $^{a}$ | 63.9% $^{a}$ | 93.4 % $^{d}$ |
| HM006 | 46.4% $^{ab}$ | 25.2% $^{ab}$ | 58.6 % $^{abcd}$ |
| KH35c | 68.3% $^{a}$ | 28.4% $^{abc}$ | 89.3 % $^{d}$ |
| KH46 | 68.9% $^{a}$ | 7.0% $^{bc}$ | 90.0 % $^{d}$ |
| M270 | 37.5% $^{abcd}$ | 6.8% $^{bc}$ | 37.0 % $^{abc}$ |
| 2011 | 15.5% $^{bcd}$ | 8.6% $^{bc}$ | 84.8 % $^{d}$ |
| Rm41 | 13.9% $^{bcd}$ | 1.7% $^{c}$ | 50.5 % $^{abcd}$ |
| RU11/001 | 39.4% $^{abcd}$ | 30.0% $^{ab}$ | 89.0 % $^{d}$ |
| SM11 | 63.4% $^{a}$ | 29.1% $^{ab}$ | 86.6 % $^{d}$ |
| T073 | 0.4% $^{d}$ | 1.8% $^{c}$ | 0.00 % $^{b}$ |
| USDA 1157 | 8.3% $^{cd}$ | 9.1% $^{bc}$ | 19.7 % $^{ab}$ |

**Table S3**. Linear regression models for the three competition experiments, performed by PhenotypeSeeker with 3-fold train/test splits of samples. The averaged model evaluation metrics of both training and test set are reported.

| Dataset | | The mean squared error | The coefficient of determination ($R^2$) | The Pearson correlation and p-value | The Spearman correlation coefficient and p-value |
|---|---|---|---|---|---|
| **Vs Rm1021** | Training set | 0.01 | 0.86 | 0.92, 0.0 | 0.97, 0.0 |
| | Test set | 0.06 | -0.75 | 0.9, 0.07 | 0.97, 0.03 |
| **Vs BL225C** | Training set | 0.01 | 0.72 | 0.85, 0.01 | 0.9, 0.0 |
| | Test set | 0.01 | 0.71 | 0.85, 0.11 | 0.87, 0.09 |
| **Vs AK83** | Training set | 0.0 | 0.99 | 0.99, 0.0 | 0.95, 0.0 |
| | Test set | 0.01 | 0.78 | 0.92, 0.05 | 0.93, 0.06 |

**Table S6 -** List of COG classes.

| COG ID | COG name |
|---|---|
| J | Translation, ribosomal structure and biogenesis |
| A | RNA processing and modification |
| K | Transcription |
| L | Replication, recombination and repair |
| B | Chromatin structure and dynamics |
| D | Cell cycle control, cell division, chromosome partitioning |
| Y | Nuclear structure |
| V | Defense mechanisms |
| T | Signal transduction mechanisms |
| M | Cell wall/membrane/envelope biogenesis |
| N | Cell motility |
| Z | Cytoskeleton |
| W | Extracellular structures |
| U | Intracellular trafficking, secretion , and vesicular transport |
| O | Post-translational modification, protein turnover, chaperones |
| X | Mobilome: prophages, transposons |
| C | Energy production and conversion |
| G | Carbohydrate transport and metabolism |
| E | Amino acid transport and metabolism |
| F | Nucleotide transport and metabolism |
| H | Coenzyme transport and metabolism |
| I | Lipid transport and metabolism |
| P | Inorganic ion transport and metabolism |
| Q | Secondary metabolism biosynthesis, transport and catabolism |
| R | General function prediction only |
| S | Function unknown |

**Table S8**. Strains and plasmids used in this work.

| Species | Strains (or plasmids) | Source/Description | Resistances | Reference |
|---|---|---|---|---|
| *Sinorhizobium meliloti* | AK83 | | | (12) |
| | 1021 | SU47 *str*-21 | Str[1] | (14) |
| | BL225C | | | (15) |
| | KH46 | | | (16) |
| | CCMM B554 | | | (19) |
| | T073 | | | (16) |
| | Rm41 | | | (21) |
| | HM006 | | | (16) |
| | GR4 | | | (22) |
| | 2011 | SU47 | Str | - |
| | USDA1157 | | | (17) |
| | KH35c | | | (16) |
| | SM11 | | | (25) |
| | RU11/001 | | Str | (27) |
| | M270 | | | (16) |
| | AK58 | | | (12) |
| | BM685 | AK83 pBHR - mRFP. | Rif[2] & Tc[3] | (29) |
| | BM687 | 1021 pBHR - mRFP. | Str & Tc | (29) |
| | GE0346 | BL225C pBHR – mRFP. | Rif & Tc | This work |
| | GE0323 | KH46 pHC60 | Rif & Tc | This work |

| | | | | |
|---|---|---|---|---|
| | GE0326 | CCMM B554 pHC60 | Rif & Tc | This work |
| | GE0327 | T073 pHC60 | Rif & Tc | This work |
| | GE0328 | Rm41 pHC60 | Rif & Tc | This work |
| | GE0329 | HM006 pHC60 | Rif & Tc | This work |
| | GE0330 | GR4 pHC60 | Rif & Tc | This work |
| | GE0339 | 2011 pHC60 | Str & Tc | This work |
| | GE0341 | USDA1157 pHC60 | Rif & Tc | This work |
| | GE0342 | KH35c pHC60 | Rif & Tc | This work |
| | GE0345 | SM11 pHC60 | Rif & Tc | This work |
| | GE0357 | RU11/001 pHC60 | Rif & Tc | This work |
| | GE0359 | M270 pHC60 | Rif & Tc | This work |
| | GE0360 | AK58 pHC60 | Rif & Tc | This work |
| *Escherichia coli* | BM266 | S17-1 λpir pHC60 | Tc | (29) |
| | BM679 | S17-1 λpir pBHR- mRFP | Tc | (29) |
| Plasmids | pBHR-mRFP | Constitutive expression of RFP | Tc | (2) |
| | pHC60 | Constitutive expression of GFP | Tc | (1) |

### Supplemental material references


1. Cheng HP, Walker GC. 1998. Succinoglycan is required for initiation and elongation of infection threads during nodulation of alfalfa by Rhizobium meliloti. J Bacteriol 180:5183–5191.

2. Smit P, Raedts J, Portyanko V, Debellé F, Gough C, Bisseling T, Geurts R. 2005. NSP1 of the GRAS protein family is essential for rhizobial Nod factor-induced transcription. Science (80- ) 308:1789–1791.

3. Pini F, Spini G, Galardini M, Bazzicalupo M, Benedetti A, Chiancianesi M, Florio A, Lagomarsino A, Migliore M, Mocali S, Mengoni A. 2014. Molecular phylogeny of the nickel-resistance gene nreB and functional role in the nickel sensitive symbiotic nitrogen fixing bacterium Sinorhizobium meliloti. Plant Soil 377:189–201.

4. Ramachandran VK, East AK, Karunakaran R, Downie JA, Poole PS. 2011. Adaptation of Rhizobium leguminosarum to pea, alfalfa and sugar beet rhizospheres investigated by comparative transcriptomics. Genome Biol 12:R106.

5. R Development Core Team. 2011. R: A language and environment for statistical computing. R Found Stat Comput.

6. Seemann T. 2014. Prokka: Rapid prokaryotic genome annotation. Bioinformatics 30:2068–2069.

7. Page AJ, Cummins CA, Hunt M, Wong VK, Reuter S, Holden MTG, Fookes M, Falush D, Keane JA, Parkhill J. 2015. Roary: Rapid large-scale prokaryote pan genome analysis. Bioinformatics 31:3691–3693.

8. Tamura K, Nei M, Kumar S. 2004. Prospects for inferring very large phylogenies by using the neighbor-joining method. Proc Natl Acad Sci U S A 101:11030–11035.

9. Kumar S, Stecher G, Li M, Knyaz C, Tamura K. 2018. MEGA X: Molecular evolutionary genetics analysis across computing platforms. Mol Biol Evol 35:1547–1549.

10. Pagès H, Aboyoun P, Gentleman R, DebRoy S. 2017. Biostrings: Efficient manipulation of biological strings. R Packag version 2460.

11. Galardini M, Mengoni A, Brilli M, Pini F, Fioravanti A, Lucas S, Lapidus A, Cheng J, Goodwin L, Pitluck S, Land M, Hauser L, Woike T, Mikhailova N, Ivanova N, Daligault H, Bruce D, Detter C, Tapia R, Han C, Teshima H, Mocali S, Bazzicalupo M, Biondi EG. 2011. Exploring the symbiotic pangenome of the nitrogen-fixing bacterium Sinorhizobium meliloti. BMC Genomics 12:235.

12. Giuntini E, Mengoni A, De Filippo C, Cavalieri D, Aubin-Horth N, Landry CR, Becker A, Bazzicalupo M. 2005. Large-scale genetic variation of the symbiosis-required megaplasmid pSymA revealed by comparative genomic analysis of Sinorhizobium meliloti natural strains. BMC Genomics2005/11/15. 6:158.

13. Galibert F, Finan TM, Long SR, Puhler A, Abola P, Ampe F, Barloy-Hubler F, Barnett MJ, Becker A, Boistard P, Bothe G, Boutry M, Bowser L, Buhrmester J, Cadieu E, Capela D, Chain P, Cowie A, Davis RW, Dreano S, Federspiel NA, Fisher RF, Gloux S, Godrie T, Goffeau A, Golding B, Gouzy J, Gurjal M, Hernandez-Lucas I, Hong A, Huizar L, Hyman RW, Jones T, Kahn D, Kahn ML, Kalman S, Keating DH, Kiss E, Komp C, Lelaure V, Masuy D, Palm C, Peck MC, Pohl TM, Portetelle D, Purnelle B, Ramsperger U, Surzycki R, Thebault P, Vandenbol M, Vorholter FJ, Weidner S, Wells DH, Wong K, Yeh KC, Batut J. 2001. The composite genome of the legume symbiont Sinorhizobium meliloti. Science (80- ) 293:668–672.

14. Meade HM, Long SR, Ruvkun GB, Brown SE, Ausubel FM. 1982. Physical and genetic characterization of symbiotic and auxotrophic mutants of Rhizobium meliloti induced by transposon Tn5 mutagenesis. J Bacteriol1982/01/01. 149:114–122.

15. Carelli M, Gnocchi S, Fancelli S, Mengoni A, Paffetti D, Scotti C, Bazzicalupo M. 2000. Genetic diversity and dynamics of Sinorhizobium meliloti populations nodulating different alfalfa cultivars in Italian soils. Appl Environ Microbiol 66:4785–4789.

16. Sugawara M, Epstein B, Badgley BD, Unno T, Xu L, Reese J, Gyaneshwar P, Denny R, Mudge J, Bharti AK, Farmer AD, May GD, Woodward JE, Medigue C, Vallenet D, Lajus A, Rouy Z, Martinez-Vaz B, Tiffin P, Young

ND, Sadowsky MJ. 2013. Comparative genomics of the core and accessory genomes of 48 Sinorhizobium strains comprising five genospecies. Genome Biol2013/02/22. 14:R17.
